# Alterations in Dynamic Spontaneous Network Microstates in Mild Traumatic Brain Injury: A MEG Beamformed Dynamic Connectivity Analysis

**DOI:** 10.1101/596155

**Authors:** Marios Antonakakis, Stavros I. Dimitriadis, Michalis Zervakis, Andrew C. Papanicolaou, George Zouridakis

## Abstract

Dynamic functional connectivity (DFC) analysis has attracted interest in the last years for the characterization of brain electrophysiological activity at rest. In this work, we investigated changes in mild Traumatic Brain Injury (mTBI) patients using magnetoencephalographic (MEG) resting-state recordings and a DFC approach. The activity of several well-known brain rhythms was first beamformed using linearly constrained minimum norm variance of the MEG data to determine ninety anatomical brain regions of interest. A DFC graph was formulated using the imaginary part of phase lag value which were obtained from 30 mTBI patients and 50 normal controls. Filtering each quasi-static graph statistically and topologically, we estimated a normalized Laplacian transformation of every single quasistatic graph based on the degree of each node. Then, the related eigenvalues of the synchronization of each node were computed by incorporating the complete topology. Using the neural-gas algorithm, we modelled the evolution of the eigenvalues for each group, resulting in distinct FC microstates (FCμstates). Using the so-called *chronnectomics* (transition rate, occupancy time of FCμstate, and Dwell time) and complexity index over the evolution of the FCμstates, we evaluated the level of discrimination and derived statistical differences between the two groups. In both groups, we detected equal number of FCμstates with statistically significant transitions in the δ, α, β, and γ_low_ frequency bands. The discrimination rate between the two groups was very high in the θ and γ_low_ bands, followed by a statistically significant difference between the two groups in all the chronnectomics and the complexity index. Statistically significant differences in the degree of several anatomical subnetworks (BAN – brain anatomical networks: default mode network; frontoparietal; occipital; cingulo-opercular; and sensorimotor) were revealed in most FCμstates for the θ, α, β, and γ_low_ brain rhythms, indicating a higher level of communication within and between the BAN in the mTBI group. In our previous studies, we focused on intra- and inter-frequency couplings of static FC. Our current study summarizes a complete set of frequency-dependent connectomic markers of mTBI-caused alterations in brain connectivity that potentially could also serve as markers to assess the return of an injured subject back to normality.

## 1. Introduction

Mild traumatic brain injury (mTBI) seems to include approximately 90% from most of the brain injuries (Len and Neary, 2011), establishing it as a major root of brain insult (Huang et al., 2014). A considerable part of these patients develops persistent cognitive deficits (van der Naalt et al. 1999; Vanderploeg et al. 2005), and post-concussion symptoms can cause irremediable problems in approximately 20% of the patients (Bharath et al., 2015) several months after the first injury (Huang et al., 2014). The main characteristics of those symptoms are often physical, emotional, cognitive, and sleep disturbances and may need several months to recover (Huang et al., 2014). In many neuropsychological studies, it has reported a reduced cognitive efficiency in several brain functions, especially in tests measuring executive function, processing speed, attention, connectivity, and memory in mTBI patients with persistent symptoms (Huang et al., 2014; Pang et al., 2016). Handling mTBI patients is not a trivial task and it is most of the times affects crucially the brain functionality (Vanderploeg et al., 2005; De Monte et al., 2006).

The conventional structural neuroimaging (computed tomography (CT), acute magnetic source imaging (MRI), or functional MRI (fMRI)) usually offers low sensitivity profiles on the detection of physiological alterations caused by mTBI (Kirkwood et al., 2006). Magnetoencephalography (MEG) is a non-invasive functional modality that detects activity from the synchronous oscillations of neurons’ membranes in the gray matter. Thus, MEG is a modality that mainly incorporates high sensitivity keeping the environmental noise to a low level, and includes low resolution spatial details and highly temporal accuracy (Leahy et al., 1998). In this study, we combined for the very first time the reconstructed source MEG activity with the notion of Functional Connectivity (FC) for the characterization of mTBI over the evolution of time. FC is crucial for the characterization most of the brain disorders (Eierud et al., 2014). The term FC was coined when the human brain was first modelled as a neurophysiological network with functional communication among several anatomical areas. These distinct networks can exist in a range of spatiotemporal scales with spatial diversity and temporal variability. Spatially, they can vary between microscopic neuronal level and large-scale interconnected areas (Eierud et al., 2014).

Several studies, including ours, recently investigated the design of the robust biomarker for the identification of mTBI including MEG and FC (see reviews by Jeter et al., 2013; Huang et al., 2009; and Eierud et al., 2014); Castellanos et al., 2010; Zouridakis et al. 2012; 2013; Da Costa et al. 2015; Vakorin et al. 2016 Dunkley et al. 2015). More recently, Dunkley et al. (2018) investigated the impact of injury underlying ICNs and showed increased coupling in the default mode network of mTBI patients. Dimitriadis et al. (2015b) using phase-synchronization quantified intra-frequency couplings at the sensor level and showed that significantly different patterns were seen mostly in the delta band, whereas Alhourani et al. (2016) used the same metric at the source level and showed reduced local efficiency in different brain regions in mTBI patients. In a follow-up study, Antonakakis and colleagues (2015, 2016, 2017a) showed less dense for mTBIs in which was in line with (Rapp et al., 2015), and later (Antonakakis et al., 2017b) that mTBIs obtained a higher synchronization among rich-club hubs. More recently, Li et al. (2018) showed a denser causality network for mTBI patients, whereas Kaltiainen et al. (2018) underlined that aberrant theta-band activity can provide an early objective sign of brain abnormality after mTBI.

MEG-based FC is noticed as an emerging prcedure in the development of reliable biomarkers for mTBI using resting-state networks (RSNs), not only at the sensor level, but also at source level, since such RSNs have been successfully estimated in past few years (Brookes et al., 2011a,b; Luckhoo et al., 2012; Hipp et al., 2012; Hall et al., 2013; Wens et al., 2015). MEG-based RSNs seem a promising approach for detecting several other brain functional abnormalities, involving dyslexia (Dimitriadis et al., 2013b, 2015c; 2016c), mild cognitive impairment (Dimitriadis et al., 2015b) and multiple sclerosis (Tewarie et al., 2015). However, summarizing RSNs as short-lived transient brain states varies significantly across different studies, ranging from estimating the MEG frequency spectrum (Vidaurre et al., 2016) to band-specific amplitude envelopes of reconstructed MEG sources (Baker et al., 2014; O’Neill et al., 2015 Vidaurre et al. 2016). A recent neuroimaging index, called chronnectomics (Allen et al., 2012; Calhoun et al., 2014), was showed in fMRI studies and it combines the synergy between the time-varying FC and the evolution of the distinct spatio-temporal alternations among the brain states (Dimitriadis et al., 2013a; Dimitriadis et al., 2015a). To our knowledge, this is the first study that utilizes DFC on the reconstructed source activity from MEG for the investigation of mTBI. Similar methodological aspects are adopted by a recent study (Dimitriadis et al., 2018a).

In this study, we investigated whole-brain dynamic FC (DFC) derived from MEG resting-state data from 80 subjects. The within brain interactions were modeled using beamformed MEG source activation and DFC among ninety atlas-based brain areas using a template MRI. Each quasi-static FC was determined by the imaginary content of the phase locking value (iPLV) and it was filtered statistically and topologically (Dimitriadis et al., 2017a,b) for the reduction of spurious connections. Subsequently, we coded the estimated time-varying network activity into prototypical network microstates or FCμstates (Dimitriadis et al., 2013a, 2018a). Through this approach, we derived symbolic-formed time series for which the chronnectomic behavior (Dimitriadis et al., 2018a) was modeled based on the metric transition rate of FCμstates, fractional occupancy of each FCμstate, Dwell time and complexity (Antonakakis et al., 2016b; Dimitriadis, 2018b). Finally, we assessed differentiations in the DFC of mTBI using statistical inference and classification.

## 2. Methods

### 2.1. Participants and Recordings

The current study included fifty right-handed healthy controls (HC) (29.25 ± 9.1 years of age) and thirty right-handed mTBI patients (29.33 ± 9.2 years of age) (Gouridakis et al., 2012). All clinical information including selection criterions and patients’ demographics were reviewed and provided by certified clinicians and they are available in our previous studies (Gouridakis et al., 2012; Antonakakis et al., 2016; 2017a,b). The HC group was determined at UTHSC-Houston and the recruitment of the mTBI patients was done by three trauma centers in the greater Houston metropolitan area. Those centers participated in larger study and detailed information can be found elsewhere (Gouridakis et al., 2012). All the participants had given their written and informed consent to participate in the current study. Further details are provided in our previous studies (Gouridakis et al., 2012; Dimitriadis et al., 2015b; Antonakakis et al., 2016; 2017a,b). The definition of the every mTBI patient was based on the guidelines the American Congress of Rehabilitation Medicine (Kay et al., 1993) and of department of Defense (Assistant Secretary, 2007). The corresponding approval for the project was declared by the Institutional Review Boards (IRBs) at the participating institutions and the Human Research Protection Official’s review of research protocols for the department of Defense. All procedures were fully compliant with the Health Insurance Portability and Accountability Act (HIPAA).

The MEG activity was acquired with a whole-head Magnes WH3600 system of 248 axial gradiometers (4D Neuroimaging Inc., San Diego, CA) for ten minutes. Every subject was in a supine position with eyes closed during the acquisition of spontaneous activity. The sampling rate was 1017.25 Hz and an online bandpass filter between 0.1–200 Hz was applied. No independent ocular and cardiac activity monitoring was recorded. In the current study, we analyzed the five minutes out of ten minutes after excluding activity contaminated with artifacts (Dimitriadis et al. 2015a) and the conversion from axial gradiometer recordings to planar gradiometer field approximation in FieldTrip (Oostenveld et al., 2011).

### 2.2 MEG Preprocessing

The artifact reduction in the MEG recordings was performed within an automated detection and elimination procedure as it was described elsewhere (Antonakakis et al., 2017a) including the software FieldTrip (Oostenveld et al., 2011) and implemented routines in MATLAB (The MathWorks, Inc., Natick, MA, USA). In brief, the noisy activity was reduced including the following steps: (1) correction of the bad MEG channel activity applying interpolation techniques (2) frequency spectrum elimination within the range 0 – 100 Hz using digital filtering (3) restricted filtering of the power line noise at 60 Hz with a notch filter and (4) detection and elimination of the biological artifacts (ocular and cardiac) by the decomposition of the MEG data into statistically independent components (Delorme and Makeig, 2004) and the application of the combined fixed thresholds on the metrics, kurtosis, skewness and Rényi entropy (Antonakakis et al., 2017a).

### 2.3 Source Analysis

Atlas-based beamforming was applied reconstructed the source activity from the MEG data for every well-known brain rhythm. The investigated frequency content included δ (0.5–4 Hz), θ (4–8 Hz), α_low_ (8–10 Hz), α_high_ (10–13 Hz), β (13–30 Hz), γ_low_ (30–55 Hz), and γ_high_ (55–90 Hz). First, the MEG sensors of every subject were realigned and normalized using a standard template T1 weighted MRI of 2 mm resolution as it was provided by SPM8 (Weiskopf et al., 2011). The clustering of the MRI voxels into ninety brain regions of interest (ROIs) was performed based on the Automated Anatomical Labeling (AAL) atlas (Tzourio-Mazoyer et al., 2002; Hillebrand and Barnes, 2002; Hillebrand et al., 2016; Hunt et al., 2016; Dimitriadis et al., 2018a). Employing a realistically shaped head model of one single shell (Nolte et al., 2003), we included 5061 sources (6 mm resolution) that covered the entire brain tissue. The frequency-depended MEG source activity was reconstructed using the linearly constrained minimum norm variance (LCMV) algorithm in FieldTrip.

We adopted similar methodological modules as in a recent study (Dimitriadis et al., 2018a) for the determination of a representative source signal in every ROI. In particular, the contribution of every MEG sensors was weighted by the LCMV beamformer for the reconstruction of the voxel-based time series for the entire predefined grid. The projection of the MEG sensor activity to the source point was the definition of the each spatial filter. Each atlas-based ROI contained a different number of voxels. Subsequently, we estimated the ROI-representative by functionally interpolating activity from the voxel time series of the specific ROI. Within each atlas ROI, we estimated the correlation between every pair of source time series creating a functional connectivity representation. The next step was the calculation of the node-strength of each voxel within the ROI. The strength was determined by summing the existed connectivity values between the specific node-voxel with the other node-voxels within the same ROI. Then, we normalized every strength resulting to a set of weights with sum equal to 1 within the ROI. The procedure ended up with the estimation of a representative time series for each ROI by getting the sum across the multiplying voxel time series with their respective weights. A block diagram of the procedure is given (Fig.1a) with exemplary a time series from a specific ROI.

**Figure 1.**
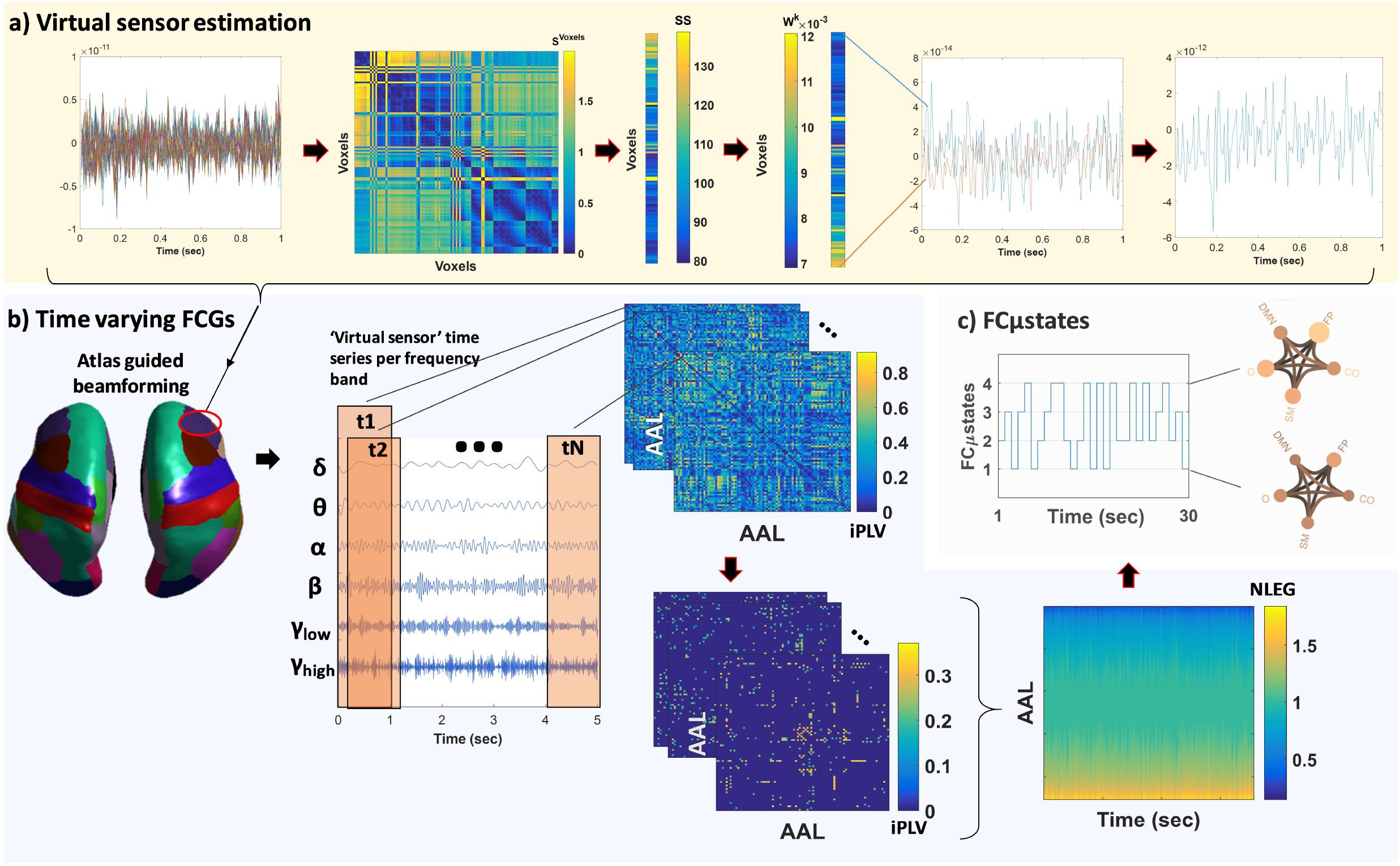
A brief overview of the proposed pipeline. **a)** Steps for determining the representative virtual sensor for each ROI. (Left to right) A sample of the 119 voxel time series from the right inferior frontal gyrus. Distance correlation matrix (S^Voxels^) derived by pair-wise estimation of the 119 voxel time series. Summation of the columns of S^Voxel^ produced the vector SS. Normalization of vector SS further producesd W^k^ where its sum equals to 1. Multiplication of every voxel time series with the related weight from the W^k^. This example demonstrates multiplication for the first and last voxel time series. Summing the weighted versions of every voxel time series for the final estimated virtual sensor for each ROI **b)** Basic steps for estimating time varying (or dynamic) functional connectivity graphs (DFCGs) (filtering the virtual sensor in several frequency bands given from a ROI; topologies of snapshots of imaginary part of the phase locking value (iPLV) DFGC from the first two (t1 and t2) and the last (tN) temporal segments in the δ frequency band for Subject 1. These topologies are statistically and topologically filtered; Dynamic evolution of the eigenvalues of the normalized Laplacian matrices (NLEG) for each frequency band. An example for the δ frequency band **c)** Short representation of the FCμstates procedure showing a sample of the symbolic time series and the corresponding prototypical FCμstates in a circular visualization with nodes the degree of basic anatomical networks (default mode network – DMN; frontoparietal – FP; occipital – OCC; cingulo-percular – CO; sensorymotor – SM)) and edges the strength of the iPLV. An example with the first and forth FCμstates for the 6 frequency band of Subject 1.

### 2.4 Dynamic Functional Connectivity Graphs

In the current work, DFC graphs (DFCG) were calculated and examined for each of the previously described brain rhythm. The used FC estimator was the iPLV (i.e. the imaginary side of the Phase Locking Value (Lachaux et al., 1999), a metric that has shown good sensitivity to non-zero-phase lags and tolerability to instantaneous self-interactions from volume conductance (Nolte et al., 2004). Here, we applied the specific metric in a dynamic manner for understanding better the time-varying alterations of the phase-to-phase interactions. This was achieved by computing the iPLV within a series of shifted and overlapped windows spanning the entire five minute continuous ROI time series (Fig.1b). The window-length of each temporal segment (or timestamp) was set to 2 sec with an overlap of 20% for each frequency band. The resulting number of timestamps was equal to 1785 per frequency and subject.

#### 2.4.1 DFCG Filtering

##### Statistical Filtering of the DFCG

A surrogate analysis was performed to evaluate the non-spurious iPLV connections on each sliding window (i.e., timestamp) for every specific brain rhythm. Our examined null hypothesis H_0_ had to do with whether the given iPLV coupling belongs to the empirical distribution estimated by the surrogates. To evaluate this hypothesis, ten thousand surrogate time-series were generated. Each surrogate quasi-static FCG was defined by shuffling temporally the original entire representative ROI time series (Aru et al., 2015). This procedure ensured similar statistical properties for both original and surrogate iPLV (iPLV_s_). After the estimation of the empirical distribution, a statistical level of significance was determined for every iPLV value. The determination of this level was done by estimating the amount of iPLV_s_ that was higher than the original iPLV. The *p*-value was ste to 0.05. A further condition was applied for examination of multiple comparisons within each quasi-static FCG (a 90 × 90 matrix with tabulated p-values) with the expected fraction of false positives being at the level 0.01 (Benjamini and Hochberg, 1995). The non-significant values were set to zero and the final DFCG had a 3D dimension of 1785 (segments) × 90 (sources) × 90 (sources) per subject and brain rhythm.

##### Topological Filtering of the DFCG

In addition to statistical filtering, a data-based topological connection-cutting was applied based on a recently suggested procedure (Dimitriadis et al., 2017a, b). The so-called Orthogonal Minimal Spanning Trees (OMST^1^) was performed to uncover the entire structure of the most dominant paths within every quasi-static FCG. The the OMST procedure was first enhanced from the notion that the full connected FCG can be extracted to an acyclic FCG or MST of minimum cost from the root node to leaf node without changing the strong connections strong from the weak connections (Dimitriadis et al., 2017a, b). Subsequently, the global efficiency of the specific MST was optimized preserving same total cost among within the connections (Dimitriadis et al., 2017b, d). The resulting DFC profiles had a 3D array of size 1785 (segments) × 90 (sources) × 90 (sources) for every subject and frequency band.

#### 2.4.2 DFCG Metrics

In every node of the statistically and topologically quasi-static FCG, we calculated the strength and degree. In particular, the strength was defined as the total sum of the iPLV values in every node resulting to a vector of ninety values per quasi static graph, brain rhythm and subject (90 x 1785 x 6 x 80). The degree was defined as the total number of connections in every node resulting to same dimensions as the strength. Moreover, to better understand the network connectivity in the whole brain, we estimated the mean strength between couples of the following brain networks, default mode network (DMN), sensorimotor (SM), frontopatietal (FP), occipital (O), and cingulo-opercular (OC)— and the degree for each of these networks (these abbreviations were used to present the interaction among these brain networks).

#### 2.4.3 Symbolization of the DFCG

The full symbolization scheme has been described in a great detail in specific previous studies (Dimitriadis et al., 2012a,b, 2013a,b). The symbolization scheme included the representation of the DFC profiles as prototypical functional connectivity microstates (FCμstates). This procedure exploited a vector quantization procedure for the effective modeling of DFCGs (Dimitriadis et al., 2013a). For this procedure, we used the neural-gas (NG) algorithm (Martinetz et al., 1993) which has the property of converting a FC time series into to a codebook with a constant number k of prototypical FCμstates (Dimitriadis et al., 2013a, b, 2015a). Thus, DFCG conversion was modeled based on the vectorization of the upper triangular part of every quasi-static FCGs. This procedure ended up to a 2D array with the first dimension being the number of timestamps and, the second, the vectorized quasi-static FCG (Dimitriadis et al., 2013a). In the current work, we used the statistically and topologically filtered DFCGs in their inherent format, i.e., as 3D tensors (the third dimension was time). Then, Adopting a recent approach for modeling the DFCGs (Dimitriadis 2018) based on the Laplacian transformation of the filtered DFCG eigenvalues, we first calculated the normalized Laplacian matrix for every quasi-static FCG based on the equation,

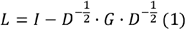

where D is the degree matrix and G is the quasi-static FCG. Afterwards, we applied eigenvalue analysis to each of the normalized Laplacian matrices. By applying the current eigenvalue analysis, we were able to reveal the synchronized level of the original FCGs. Then, the NG algorithm was applied on the 2D arrays after the concatenation across all subjects (90 x [timpestamps x subjects]) independently for each frequency band and group (control: 90 x 89250 and mTBI: 90 x 53550). By applying this, the existed richness of information in the DFC profiles was represented by a decomposed matrix U indicating the assignment of input FC profiles to code vectors.

The NG algorithm was an iterative procedure which was applied in the normalized Laplacian matrix and the stop condition was based on the threshold. The detection of the optimal threshold was obtained based on the estimation the reconstructed error between the original and the reconstructed 2D matrix (90 × [slides × subjects]). The final k was assigned to each timestamp of the DFCG based on a predefined threshold per case. Notice that the selection of parameter k expressed the compression level and fidelity.In the present study, the stop condition of the reconstruction error was < 1% for every the brain rhythm and group. The visualization of the reconstructed error E versus the threshold was used for the observation its corresponding plateau resulting to the selection of the final threshold. The estimated normalized Laplacian-based symbolic times series that preserved the information of network FCμstates are indicated hereafter as STS^L-EIGEN^ (L:Laplacian – Eigen:Eigenalysis). An STS^L-EIGEN^ can be assumed as a first order Markovian Chain that describes temporally the evolution FCμstates in both groups. The final number of FCμstates was 4 for the δ frequency band and 3 for the rest frequency bands for both groups.

#### 2.4.4 Characterization of Time-Varying Connectivity

We mainly calculated DFC metrics based on the so-called *chronnectomic* features which were estimated on the STS^L-EIGEN^ that expressed the alternation among the FCμstates. The first metric was the transition probability (TP) that expressed the transition rate among the FCμstates applying the following equation,

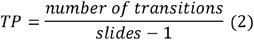

where slides denotes the number of timestamps. TP yielded higher values for increased numbers of “hops” of the brain between the FCμstates. The next metric was the Occupancy Time (OT) of the FCμstates. The OT metric accounts the percentage of the occurrence across the experimental time in every FCμstate and it is described as,

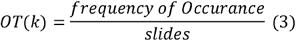

where slides denotes the number of timestamps and *k* denotes the FCμstates. The next metric was the complexity index (CI). The CI of a STS^L-EIGEN^ is estimated as follows,

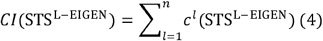

where *c^l^*(STS^L-EIGEN^) was the number of distinct substrings of STS^L-EIGEN^ of length *l* (Dimitriadis et al., 2012c; 2013a; 2016a; Antonakakis et al., 2016). The parameter *l* was set to 7 for the frequency bands and subjects. An additional applied metric was the Dwell Time which counts the time the brain spends on a FCμstate before it transitions to another brain state. The difference with OT is the search of consecutive periods that a brain sticks to a brain state, while OT measures the summation of time that the brain spends on a brain state. The final used metric was the Transition Probability Matrix. This metric counts the pairwise transitions count all possible pairwise transitions of brain states over a common codebook scanning the Markovian chain from left to right. For example, the following STS describes the temporal evolution of three brain states [1 2 2 1 2] and the pairwise transitions (PT) are equal to: PT_12_ = 2/4=0.5 and PT_21_ = 1/4=0.25. The size of the pairwise transition matrix is equal to (the number of brain states) x (the number of brain states).

For the aforementioned chronnectomic features, we assessed their significant level by adopting a surrogate data analysis. We shuffled 1000 times the subject-specific STS^L-EIGEN^ resulting in 1000 surrogate chronnectomics estimates. Then, we assigned a p-value to every subject-specific chronnectomic by comparing the original value with the 1000 surrogate values. Finally, we analyzed at the group-level only the subject-specific chronnectomic features that were statistically significant (p < 0.01). Especially for the pairwise transitions, we created a transition matrix by assigning a p-value to each pairwise transition. Subsequently, we controlled for multiple comparisons employing the false detection rate correction procedure (Benjamini and Hochberg, 1995) in the level of q ≤0.01 independently in every frequency band.

#### 2.4.5 Classification and Statistical Assessments

We assessed the discrimination of the mTBI and control groups by using as feature vector all the chronnectomic indexes (TP, OT, CI and DT) of the corresponding FCμstates via machine learning schemes. Within an iterative scheme of thousand times and using ten-fold cross-validation, we first calculated the most informative features using a Laplacian score (Xe et al., 2005) through iterative bootstrapping procedure for determining a cut-off threshold. This threshold was served a cut-off condition in the features (Dimitriadis et al., 2015a; Antonakakis et al., 2016a, b) avoiding any hazard for overfitting. The labels of the two groups were first shuffled and their Laplacian score was re-calculated. Then, we applied a threshold to the original Laplacian scores derived by the mean + 2SD (SD: standard deviation) of the thousand Laplacian scores estimated via the randomization procedure. Afterward, classification based on the k nearest neighbor algorithm (kNN; Horn and Mathias, 1990) was performed to observe the discrimination level between the two groups. The entire procedure was repeated separately for each frequency band, including all FCμstates and chronnectomic indexes per group.

Statistical analysis was performed on the feature vectors used for classification to further confirm the discrimination level between the two groups. The same statistical analysis was performed on the mean strength between pairs of the brain networks (DMN, SM, FP, O, and OC) and mean degree of each brain network to evaluate the discrimination level based on the mean value between the two groups. The statistical procedure included a normality control based on the Kolmogorv-Smirnov test and based on its result, either the use of parametric pair-wise sample t-test or non-parametric pair-wise Mann-Whitney u-test (Antonakakis et al., 2016a). The threshold for significance of the p-value was set to 95%. Control for multiple comparisons was applied thought the FDR adjustment (Benjamini and Hochberg, 1995) at a q level of less than 0.01.

## 3. Results

### 3.1. From multichannel recordings to a restricted repertoire of quasi-static FCμstates

Our analytical pipeline revealed unique phase-based synchronized functional connectivity patterns (FCμstates) that can be considered as discrete brain states: the human brain ‘jumps’ between characteristic FCμstates whose temporal evolution can hide important information. Figure 2 illustrates representative state-transitions of FCμstates, from the first HC and mTBI subjects, for every frequency band. The analysis revealed four FCμstates for δ and three for θ to γ_2_ in both groups. In addition to the representative STS^L-EIGEN^, we demonstrated brain topologies that represent the mean-degree of the nodes that constitute the default mode network, independently for each FCμstate. Figure 3 demonstrates the significant transition rates between FCμstates for every group and frequency band. FCμstates are sketched as 5-to-5 networks, where the size and color of every node codes the mean degree within every subnetwork, while the color of the between sub-networks connections decodes the functional strength between the ROIs that constitute those subnetworks.

**Figure 2.**
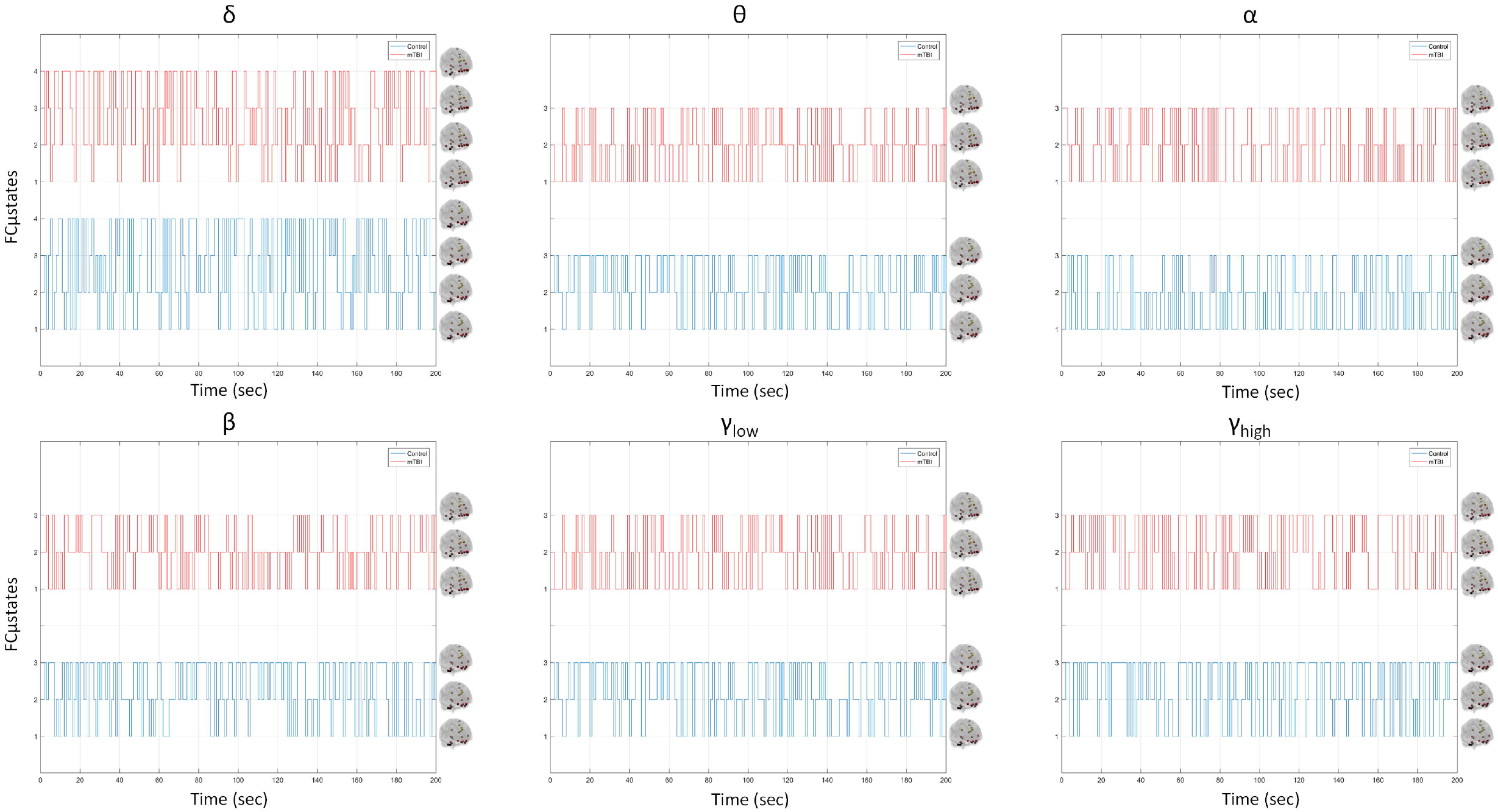
Temporal evolution of FCμstates for representative HC and mTBI subjects in every frequency band. Sample symbolic time series per frequency band for the control and mTBI group. The brain topologies represent the mean degree of every node that constitutes the default mode network for each FCμstate.

**Figure 3.**
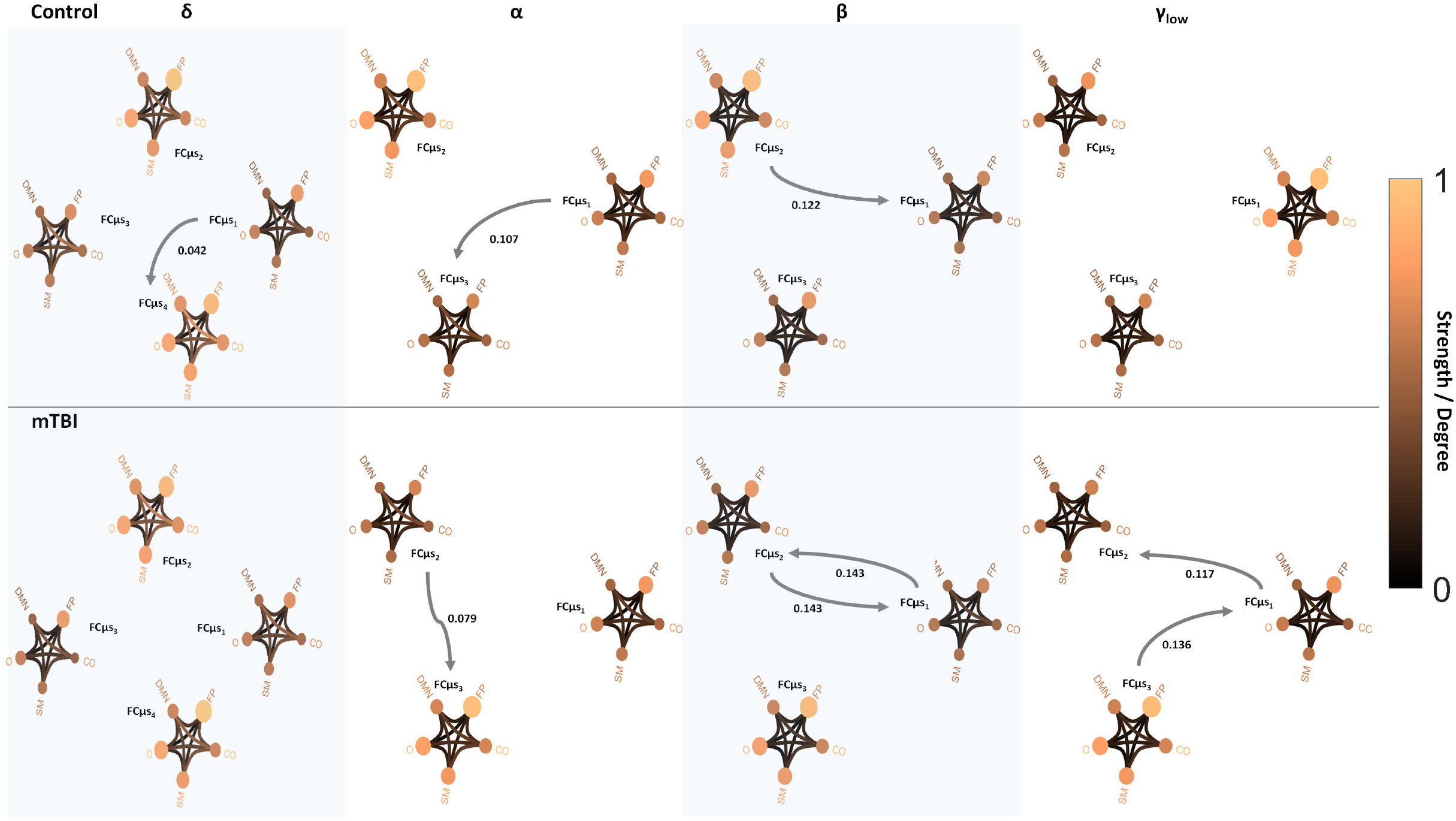
Pairwise transition diagram between FCμstates for each group and frequency band. Statistically significant group differences of the pairwise transition rates between FCμstates characteristic for every frequency band. The color and size of every node encodes the mean degree of each sub-network, while the color of between sub-networks connection encodes the mean functional strength between the ROIs that constitute each sub-network. Both strength and degree are normalized using the maximum value between the two groups and all frequency bands. The color bar is common to all cases. Grey arrows represent statistically significant pairwise transition rates between FCμstates. Every FCμstate is represented with the corresponding label FCμs_i_ (i=1,…,4).

### 3.2. Aberrant Higher-Lower Global Mean Degree for mTBI subjects

Figure 4 summarizes the global mean degree for every FCμstate across frequency bands in both groups. Following a statistical group comparison independently for every FCμstate, we revealed statistical significant differences between the groups in most cases. Our analysis under the framework of global mean degree untangled an aberrant higher-lower pattern for mTBI subjects compared to healthy controls.

**Figure 4.**
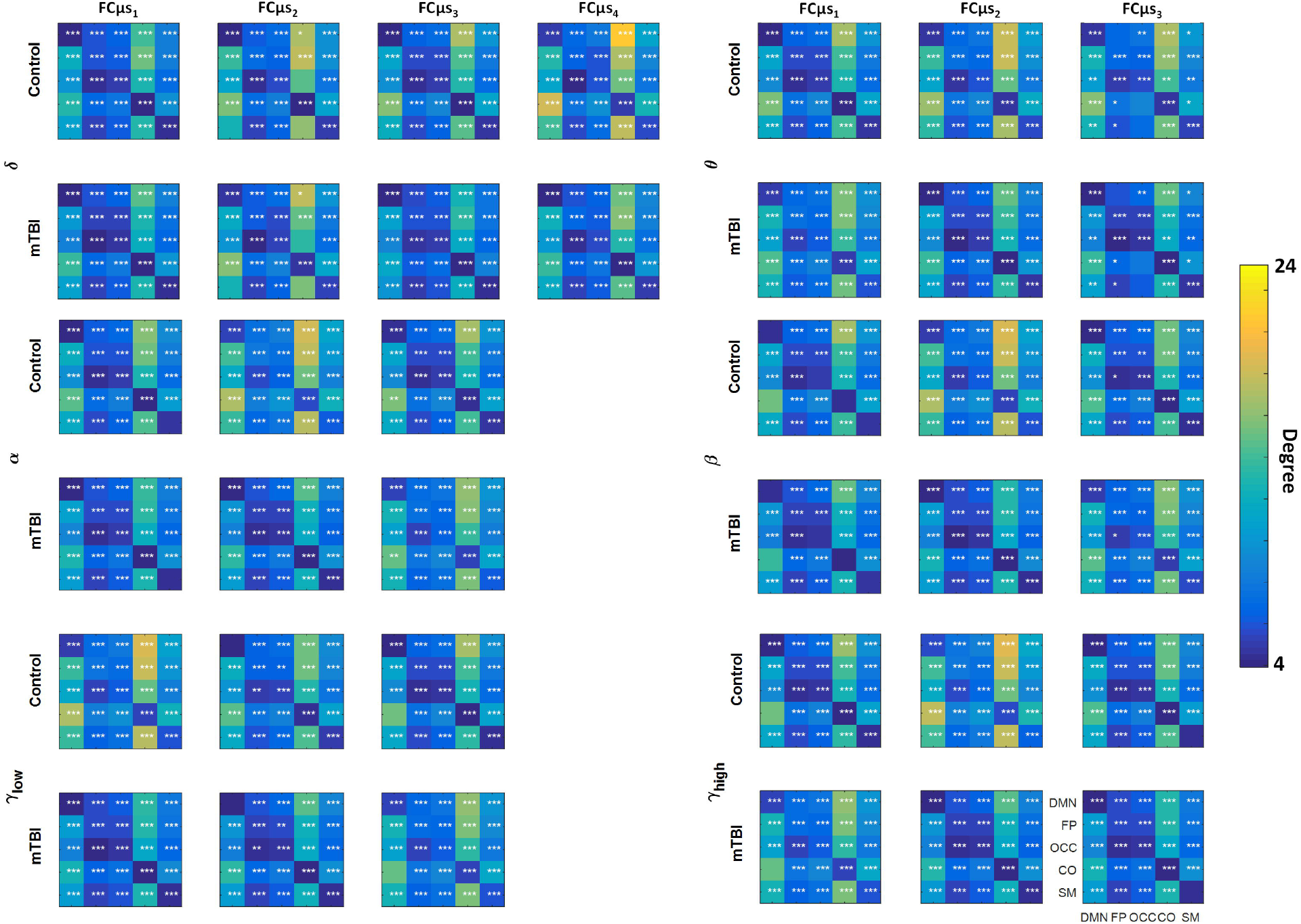
Averaged node degree. Group (control: blue and mTBI: red) averaged node degree per FCμstate (FCμs_i_ for i = 1,…,4) and frequency band (δ, θ, α, β, γ_low_, γ_high_) for every basic anatomical network (BAN: default mode network – DMN; frontoparietal – FP; occipital – OCC; cingulo-percular – CO; sensorymotor – SM)). The diagonal voxels depict averaged node degree within every BAN while the non-diagonal voxels represent averaged node degree between every pair of BANs. Statistical comparisons (*: p-value<0.05, **: p-value<0.01 and ***: p-value<0.001) are presented between the two groups. Statistical evaluation was based on the procedure described in Section 2.4.5.

### 3.3. Classification performance and Feature Ranking

Table 1.A summarizes the results of absolute classification performance based on chronnectomic features in every frequency band. Table 1.B represents the ranking of features selected consistently across the cross-validation scheme per frequency band. Figure 5a demonstrates the group-averaged chronnectomic features for every frequency band. Statistical analysis revealed significant higher TP and CI for HC compared to mTBI. The mTBI averaged OC of FCμstate was statistical significantly higher the HC subjects in the δ frequency band. Group mean values were statistical significantly higher for HC compared to mTBI for OC in the δ, θ, and α frequency bands, and higher for mTBI compared to HC in the β, and γ_low_ frequencies. Additionally, group mean DTs were significantly higher in HC compared to mTBI in the δ, θ, α, and β bands, and higher in mTBI in the γ_low_ band compared to HC. TP in the θ, β, and γ_low_ bands were the most discriminative features across the chronnectomic features. Projecting every subject using this triplet of TP-related features, absolute discrimination was obtained between the two groups, and each group was allocated a distinct projection subspace (Fig. 5.B).

**Table 1.**
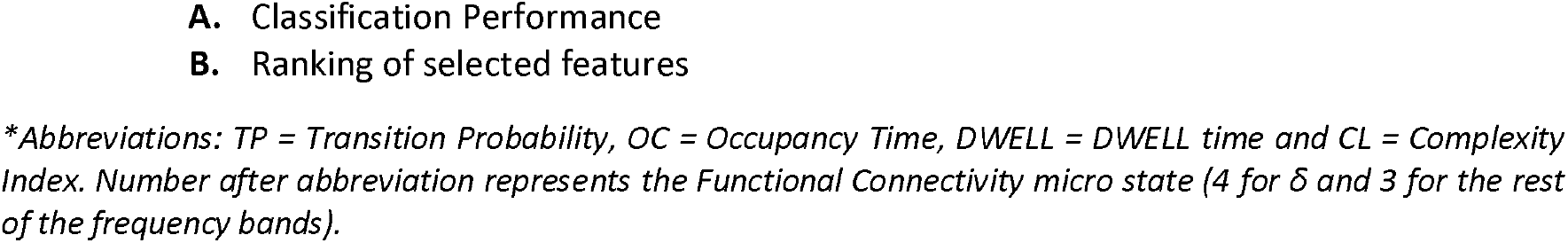
Classification performance and Feature Selection per frequency band.

**Figure 5.**
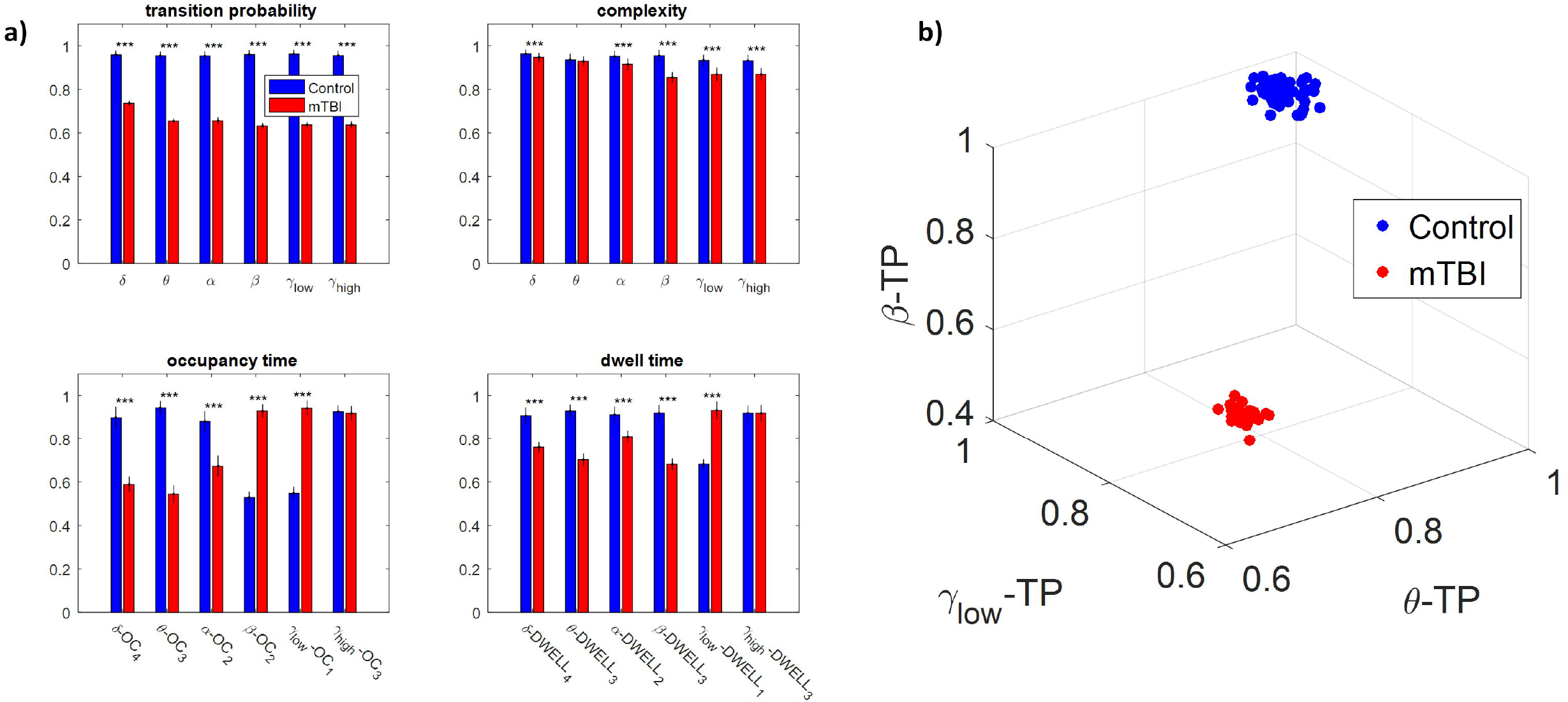
Discrimination between the two groups. **a)** Average value and standard deviation of each metric (transition probability, complexity, occupancy time, and dwell time) and group (blue for control and red for mTBI) for every frequency band. Statistically significant difference on mean value for every metric between the two groups are noticed (*: p-value<0.05, **: p-value<0.01 and ***: p-value<0.001) **b)** The most informative features, as categorized by ranking analysis, that result in complete separation between the control (blue) and the mTBI (red) groups. Statistical evaluation was based on the procedure described in Section 2.4.5.

## 4. Discussion

Thought the current study, we developed a framework for analyzing the spatiotemporal evolution of functional connectivity patterns on the MEG source-reconstructed activity at rest for mTBI and HC subjects. Each frequency-dependent DFCG was discretized via the neural-gas algorithm into a symbolic time series that described the temporal evolution of brain states (FCμstates). DFCGs were trimmed as Markovian chains of 1^st^ order from which valuable chronnectomic markers were estimated. Our results revealed significantly lower values of transition probability (TP), complexity (C), dwell time (DT), and occupancy time (OT) for mTBI subjects compared to HC. Following a machine learning approach, we succeeded to discriminate the two groups with absolute accuracy (100%). The three features with the highest discriminative power were the TP in the θ, β, and γ_low_ frequency bands.

Recent advances in MEG and network neuroscience have shown that mTBI can be manifested as an excessive pattern of slow-wave activity (Huang et al., 2012), while localization of this slow-wave activity can reveal the foci of the damage (Huang et al., 2014). Dimitriadis et al. (2015b) employed PLI to quantify time-static FCGs at the sensor level. They revealed a dense local and sparse long-range connectivity pattern for healthy controls and a sparse local and dense long-rage pattern for the mTBI subjects. Activity in the alpha frequency was the most discriminative for separating the two groups. In our previous studies (Antonakakis et al., 2015, 2016, 2017a), we employed an analysis of combined the content of intra-based and inter-based frequency couplings in a single FCG suggestion that HC formed a dense network of stronger local and global connections in agreement with other studies (Rapp et al., 2015). In our most recent study (Antonakakis et al., 2017b), the mTBI group indicated a hyper-synchronization in rich-club network organization compared to HC groups.

Structural neuroimaging combined with diffusion tensor imaging (DTI) revealed an association of mTBI with aberrant white matter microstructures (Huang et al., 2009). Specifically, a connection of focal increased slow wave activity with the location of white matter injury was shown, which was consistent with the hypothesis that the deceleration of oscillations could be caused by differentiation (Llinás et al., 1999).

A disrupted inter-regional frequency-dependent functional connectivity pattern has also been reported in combat-related blast injury using EEG (Sponheim et al., 2011). Oscillatory functional synchronization between brain areas play a pivotal role in network connectivity and support both cognition and behavior (Ward, 2003; Uhlhaas et al., 2009). The expression of this network connectivity in various frequencies at resting-state is related to the intrinsic multi-frequency organization of brain activity that is pertinent to normal brain function and dysfunction in various clinical populations (Engel et al., 2014).

The present study explored for the very first time the consequences of mTBI in human brain functionality at resting-state using MEG and DFCG analysis. Our analysis summarized DFCGs with prototypical connectivity graphs, where each graph was associated with a FCμstate (brain state). The major outcome of this analysis was a symbolic time series for the temporal description of the evolution of these brain states. Dynamic functional connectivity graphs were trimmed as Markovian chains from which a variety of chronnectomic metrics were extracted. Machine learning analysis over the derived chronnectomics produced absolute classification between healthy controls and mTBI subjects (Table 1). Ranking the features with an appropriate feature selection algorithm revealed that TP was the most discriminative feature in every frequency band. To visualize the absolute classification in 3D, we extracted the three features with the highest discriminative power, namely the TP for θ, β, and γ_low_ frequencies. Figure 5 b shows that the projections of the two groups using the three TRs occupy distinct 3D sub-spaces.

Figure 4 demonstrates the global mean degree for every FCμstate across frequency bands in both groups. Our statistical analysis applied independently to every FCμstate revealed significant differences between the two groups in most cases. Our analysis untangled an aberrant higher-lower pattern for mTBI subjects compared to healthy controls. The central brain networks more connected with the rest of the brain were the CO and DMN across all frequency bands. In mTBI subjects, DMN and CO were less connected with the rest of brain networks compared to healthy controls (Fig. 4). DMN reduced their activity during cognitive demands and may be involved to self-referential processing and internal emotional states (Power et al., 2011; Barch, 2017). Finally, CO played a central role in sustaining alertness and attention (Coste and Kleinschmidt, 2016).

Our study employed beamformed source-reconstruction of resting-state MEG activity in various frequency bands with a common MRI template for all subjects. However, a recent study explored the deviation in power and connectivity of virtual source MEG activity when using a template instead of native MRIs (Douw et al., 2017) and found that relative power, connectivity measures, and network estimates were consistent in both cases. Finally, it would be interesting to evaluate in a future study the sensitivity of the current analysis approach and chronnectomic features in detecting the return of mTBI subjects back to normality.

## 5. Conclusions

In the present study, we obtained for the very first time how mTBI affects the dynamics of functional brain networks on beamformed source-reconstructed resting-state activity. Dynamic functional graphs were trimmed as a Markovian chain via a well-established analytic framework that discretized their temporal evolution into a symbolic time series. Symbolic dynamics and chronnectomics proved valuable for the absolute discrimination of healthy controls from mTBI subjects. The TP in θ, β, and γ_low_ frequencies were the most discriminative chronnectomic features. CO and DMN were the central network hubs of the derived brain states (FCμstates), where mTBI subjects were less connected with the rest of the brain networks compared to healthy controls.

## Acknowledgement

The Department of Defense Congressionally Directed Medical Research Program W81XWH-08-2-0135 was supported the current work. This study belong to the project Integrated Clinical Protocol, conducted by the Investigators and staff of The Mission Connect Mild Traumatic Brain Injury Translational Research Consortium. The author SID was funded by an UK-EU fellowship and the MRC grant (MR/K004360/1: Behavioral and Neurophysiological Effects of Schizophrenia Risk Genes: A Multi-locus, Pathway Based Approach).

