## Supplementary material for "Alterations in Dynamic Spontaneous Network Microstates in Mild Traumatic Brain Injury: A MEG Beamformed Dynamic Connectivity Analysis": Table I

### Tables

**Table 1**. Classification performance and Feature Selection per frequency band.

A. Classification Performance

| Frequency band | Accuracy (%) | Sensitivity (%) | Specificity (%) |
| --- | --- | --- | --- |
| δ | 100±0 | 100±0 | 100±0 |
| θ | 100±0 | 100±0 | 100±0 |
| α | 100±0 | 100±0 | 100±0 |
| β | 100±0 | 100±0 | 100±0 |
| γ_low_ | 100±0 | 100±0 | 100±0 |
| γ_high_ | 100±0 | 100±0 | 100±0 |

B. Ranking of selected features

| Frequency band | 1^st^ | 2^nd^ | 3^rd^ | 4^th^ | 5^th^ | 6^th^ | 7^th^ | 8^th^ | 9^th^ | 10^th^ |
| --- | --- | --- | --- | --- | --- | --- | --- | --- | --- | --- |
| δ | TP | OC_4 | OC_2 | DWELL_4 | DWELL_2 | OC_1 | OC_3 | CL | DWELL_1 | DWELL_3 |
| θ | TP | OC_3 | DWELL_3 | OC_2 | OC_1 | DWELL_2 | DWELL_1 | CL | - | - |
| α | TP | OC_2 | OC_3 | DWELL_2 | DWELL_3 | CL | OC_1 | DWELL_1 | - | - |
| β | TP | OC_2 | OC_3 | DWELL_3 | DWELL_2 | CL | OC_1 | DWELL_1 | - | - |
| γ_low_ | TP | OC_1 | OC_2 | DWELL_1 | DWELL_2 | CL | OC_3 | DWELL_3 | - | - |
| γ_high_ | TP | CL | OC_3 | DWELL_3 | OC_1 | OC_2 | DWELL_2 | DWELL_1 | - | - |

**Abbreviations: TP = Transition Probability, OC = Occupancy Time, DWELL = DWELL time and CL = Complexity Index. Number after abbreviation represents the Functional Connectivity micro state (4 for δ and 3 for the rest of the frequency bands).*
